# Effects of continuous tactile stimulation on auditory-evoked cortical responses depend on the audio-tactile phase

**DOI:** 10.1101/2022.12.05.519195

**Authors:** Xueying Fu, Lars Riecke

## Abstract

Auditory perception can benefit from stimuli in non-auditory sensory modalities, as for example in lip-reading. Compared with such visual influences, tactile influences are still poorly understood. It has been shown that single tactile pulses can enhance the perception of auditory stimuli depending on their relative timing, but whether and how such brief auditory enhancements can be stretched in time with more sustained, phase-specific periodic tactile stimulation is still unclear. To address this question, we presented tactile stimulation that fluctuated coherently and continuously at 4Hz with an auditory noise (either in-phase or anti-phase) and assessed its effect on the cortical processing and perception of an auditory signal embedded in that noise. Scalp-electroencephalography recordings revealed an enhancing effect of in-phase tactile stimulation on cortical responses phase-locked to the noise and a suppressive effect of anti-phase tactile stimulation on responses evoked by the auditory signal. Although these effects appeared to follow well-known principles of multisensory integration of discrete audio-tactile events, they were not accompanied by corresponding effects on behavioral measures of auditory signal perception. Our results indicate that continuous periodic tactile stimulation can enhance cortical processing of acoustically-induced fluctuations and mask cortical responses to an ongoing auditory signal. They further suggest that such sustained cortical effects can be insufficient for inducing sustained bottom-up auditory benefits.

## 1. Introduction

The ability to detect and listen to relevant sounds in noisy environments is an essential part of our everyday life. Perception of auditory stimuli can be supported by stimuli in other sensory modalities. For example, lip-reading can improve the neural processing and intelligibility of auditory speech in noise (Crosse et al., 2016; Park et al., 2016; Zion Golumbic et al., 2013). While the influence of visual input on auditory perception has been studied extensively, the influence of somatosensory (tactile) stimuli has remained less clear.

Several studies have shown that brief tactile stimuli (≤ 500ms) can improve auditory perception. For example, the presentation of a brief tactile vibration to fingers can increase the perceived intensity of a simultaneously presented tone in noise (Schürmann et al., 2004). A critical factor for this auditory enhancement is the synchrony of the onsets of the auditory and tactile inputs: Gillmeister and Eimer (2007) found that tactile pulses applied to a finger or hand can improve auditory tone-detection thresholds when the pulses are synchronous with the tones, compared with asynchronous tactile pulses or no tactile stimulation. Consistent results were found in a similar study using brief 250-Hz sinusoidal tactile stimuli (Wilson et al., 2009). It has been shown that the somatosensorily-induced auditory enhancement is stronger when auditory and tactile stimuli have equal or nearby frequencies in the low frequency range around 250Hz (Wilson et al., 2010), although their relative phase does not seem to have much influence in this range (Wilson et al., 2009). Another less influential factor seems to be the spatial alignment of tactile and auditory inputs. Murray et al. (2005) found that the application of tactile pulses to a hand can accelerate the detection of an auditory noise burst, regardless of whether the tactile and auditory inputs are perceived to originate from the same or different locations. Brief tactile stimuli can also modulate the perception of more complex natural sounds, such as speech (Gick, 2008; Gick & Derrick, 2009; Reed et al., 2021; Reed et al., 2019), possibly as a consequence of the auditory enhancement described above. Regarding neural correlates, human brain studies have shown that the enhancing auditory effect of tactile stimulation may emerge in auditory cortical regions, including the primary auditory cortex (Hoefer et al., 2013) and auditory association areas (Foxe et al., 2002; Murray et al., 2005). Overall these studies provide converging evidence that brief tactile inputs can improve the perception of various auditory stimuli by enhancing cortical responses to these stimuli. They further show that the onset synchrony between auditory and tactile inputs is a critical factor.

A limitation of the previous studies is that they focused on relatively brief tactile events, although auditory perception may involve longer-lasting input signals characterized by slow amplitude fluctuations, such as speech. Similar to the visual case of lip-reading, it is conceivable that slowly varying tactile input can carry useful temporal cues to continuously inform the perceptual analysis of ongoing auditory signals. However, it is still unclear whether the aforementioned results based on brief tactile stimuli generalize to more continuous tactile stimuli fluctuating over multiple seconds. Only few studies have addressed this question, e.g., Fletcher and colleagues found that the presentation of speech-shaped tactile stimuli (time-varying stimuli resembling the waveform of auditory speech) can improve the intelligibility of simultaneously presented auditory speech in noise. This was observed in both normally-hearing listeners (Cieśla et al., 2022; Fletcher et al., 2018) and cochlear implant users (Fletcher et al., 2019; Huang et al., 2017) after they were trained on the audio-tactile speech. Similar studies with untrained listeners found partially conflicting results (Cieśla et al., 2019; Guilleminot & Reichenbach, 2022; Riecke et al., 2018; Riecke et al., 2019). Investigations with more basic stimuli (tactile pulse sequences and amplitude-modulated auditory tones) also showed tactile effects on auditory perception (Timora & Budd, 2013, 2018); however, contrary to the results based on brief stimuli, this effect was found to be negative (i.e., hampering auditory perception) and independent of whether the tactile and auditory inputs fluctuated at the same or different frequencies. At the neural level, the hampering effect on auditory perception was accompanied by enhanced cortical responses to the stimulus fluctuation when auditory and tactile stimuli had equal modulation frequencies, consistent with the previous observations of somatosensorily-induced neural response enhancement. Overall, these results based on longer-lasting, fluctuating audio-tactile stimulation show some analogies to the earlier results based on brief tactile events. However, given the inconsistency of these results, it is still unclear how continuous time-varying tactile input influences the processing and perception of ongoing auditory input.

We addressed this question in the present study at the level of basic auditory processing using continuous sinusoidal audio-tactile stimuli in the context of dip listening. Dip listening refers to listeners’ ability to catch brief ‘glimpses’ of an auditory signal in noise when the noise level is momentarily reduced (Cooke, 2006; Vestergaard et al., 2011). This release from auditory masking in temporally fluctuating noise can be observed in, for instance, speech recognition, which is usually better in noise that fluctuates vs. lacks fluctuations (Bacon et al., 1998; Fullgrabe et al., 2006; Gustafsson & Arlinger, 1994). Based on the previous findings above, we hypothesized that tactile input that fluctuates synchronously with auditory noise can amplify the fluctuations of the noise in the brain and thereby facilitate dip listening. Conversely, tactile input that fluctuates in anti-phase to the auditory noise may mask the signal during the noise dips and thereby hamper dip listening.

To test this, we presented normally-hearing participants with an auditory tone embedded in ongoing fluctuating auditory noise and applied continuous tactile stimulation that fluctuated either in-phase or anti-phase with the noise, or no tactile stimulation. We measured participants’ ability to listen to the tone using an auditory target-detection task that required them to detect a temporary loudness decrease in the tone. We also measured their cortical responses to the noise and the tone, using electroencephalography (EEG) and a frequency-tagging technique as applied in previous work (Budd & Timora, 2013; Timora & Budd, 2013, 2018). Under our hypothesis, we predicted that in-phase tactile stimulation (compared with no tactile stimulation) strengthens cortical responses to the fluctuations of the auditory noise and thereby improves auditory target detection. Conversely, anti-phase tactile stimulation (compared with no tactile stimulation) should reduce cortical responses to the tone and thereby hamper target detection.

## 2. Methods

### 2.1 Participants

Twenty-five healthy university students (12 males and 13 females; age range: 19 to 28 years; mean ± standard deviation (SD): 23.9 ± 2.6 years) were recruited for the study. All participants had normal hearing (pure-tone audiometric thresholds < 25 dB SPL (sound-pressure level) for 250-8000 Hz in both ears) and reported having no skin condition, or neurological or psychiatric disorders. One female participant was excluded from analyses due to a relatively high number of artifacts in the EEG recordings (>15% of epochs), which did not change the conclusions that can be drawn from the results. All participants gave their written informed consent and received study credits or monetary vouchers for their participation. The experimental procedure was approved by the local ethics committee of the Faculty of Psychology and Neuroscience, Maastricht University (ERCPN #_233_09_02_2021).

### 2.2 Stimuli

#### 2.2.1 Auditory Stimuli

Auditory stimuli consisted of a continuous tone embedded in a continuous fluctuating noise. To allow investigating cortical responses to the tone and the noise separately, the two stimulus components were amplitude-modulated using different ‘tagging’ frequencies (see Figure 1A). The tone was a 250-Hz sinusoid with a 37-Hz sinusoidal amplitude-modulation (depth: 100%, start phase: 0°), which was presented at a fixed sound-pressure level of 41.5 dB (sound-pressure level, SPL). The noise was generated from a Gaussian distribution and bandpass-filtered into a 1-octave range centered on the tone carrier frequency (4th order Butterworth filter with zero phase shift, cutoff frequencies: 176.8 Hz and 353.6 Hz). The noise was amplitude-modulated with a 4-Hz sinusoid (depth: 80%, start phase: 0°), which was expected to facilitate dip listening (Gustafsson & Arlinger, 1994). Because the noise level at which putative audio-tactile interactions would be strongest was unknown a priori, a range of four noise levels was explored. This further allowed investigating whether the intensity of auditory input modulates putative tactile effects, which would provide an indicator for audio-tactile integration. The following specific signal-to-noise ratios (SNRs) were used: 12.8 dB, 3.4 dB, -2.7 dB, and -7.2 dB (SPL). They were chosen based on prior informal listening tests with the aim to elicit robust frequency-tagged responses (see section 2.7.2) while reducing potential range effects. Firstly, the tone was set to a clearly audible and comfortable level. Secondly, the minimum SNR was defined as the highest noise level (*N*_*max*_) at which the tone was still audible and the noise level did not elicit any discomfort. Thirdly, the maximum SNR was defined as the lowest noise level (*N*_*min*_) at which task performance (see section 2.4) still fell below the ceiling. Finally, the intermediate SNRs(*x* = 2 *and x* = 3) were interpolated using function 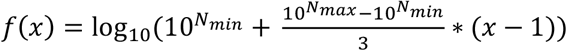 with the aim to induce linear changes in task performance.

**Figure 1.**
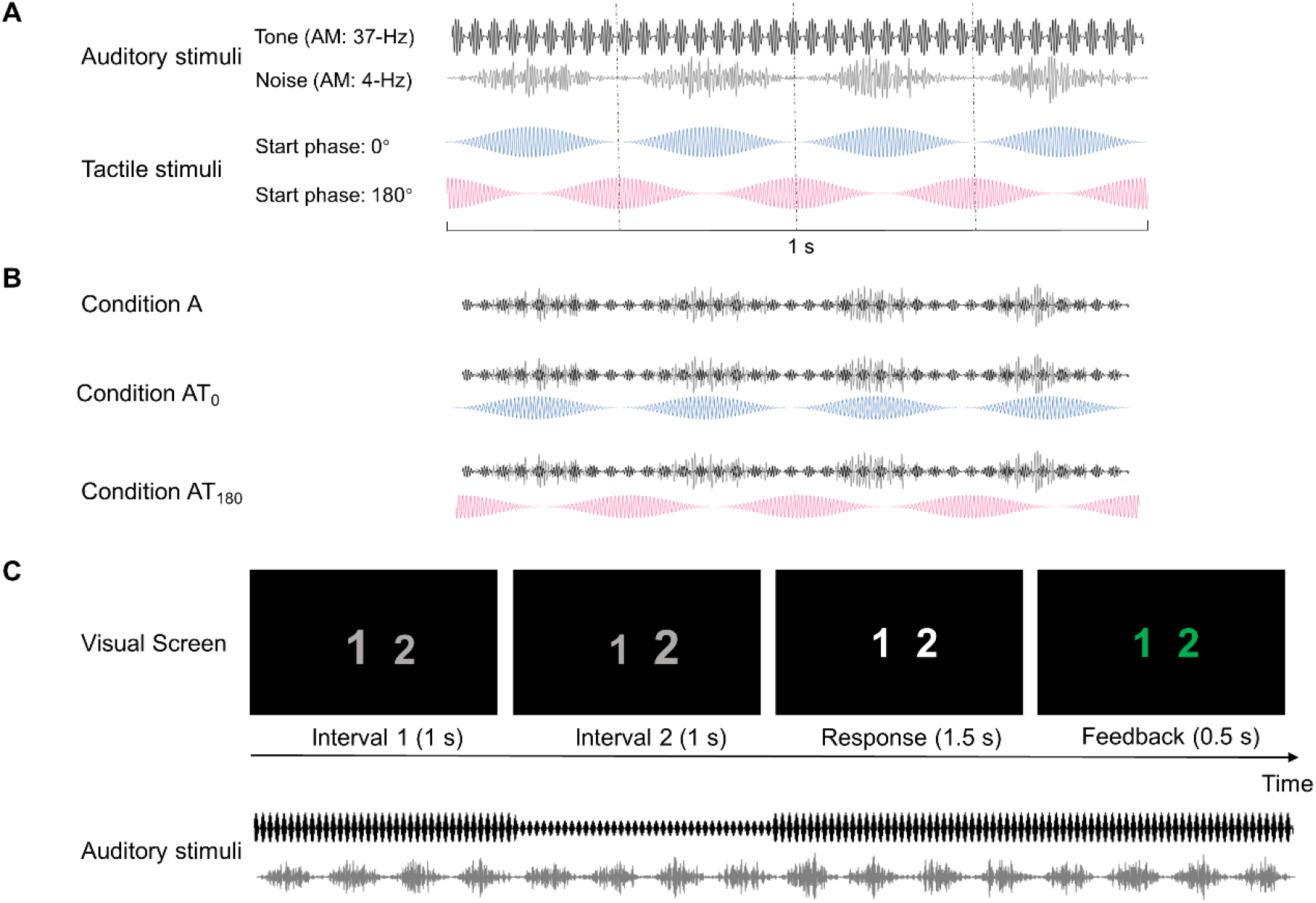
Audio-tactile stimuli and task design. **A**. Auditory and tactile stimuli. The black waveform and gray waveform illustrate the auditory tone and noise, respectively. The auditory tone carried a 37-Hz amplitude-modulation, and the auditory noise carried a 4-Hz amplitude-modulation. The other waveforms represent the tactile stimulation, which had the same amplitude-modulation as the auditory noise (4 Hz). The in-phase tactile stimulation (blue waveform) and anti-phase tactile stimulation (magenta waveform) were implemented by setting the start phase of the tactile stimulation to 0° or 180° relative to the start phase of the auditory noise. **B**. The top waveform illustrates the purely auditory condition (condition A). This condition included the 4-Hz AM noise and the 37-Hz AM tone, which were presented continuously together. The middle and bottom waveforms show respectively the in-phase audio-tactile condition (condition AT_0_) and anti-phase audio-tactile condition (condition AT_180_). These conditions were identical to condition A but additionally included the continuous 4-Hz tactile stimulation. **C**. Each trial contained two consecutive observation intervals. The digits “1” and “2” were continuously shown on a screen and consecutively increased in size to indicate the 1^st^ or 2^nd^ interval. The tone was presented throughout, with a decrease in loudness during either the 1^st^ or 2^nd^ interval (in this example during the 2^nd^ interval). Participants’ task was to detect this loudness decrease and report the interval during which it occurred by pressing a button labelled ‘1’ or ‘2’ during the response interval. Visual feedback on response correctness was presented after each trial.

#### 2.2.2 Electrotactile stimuli

Electrotactile stimulation was used because, unlike vibrotactile stimulation, it does not produce mechanical vibrations at the skin-transducer interface and therefore cannot constitute an acoustic confound in auditory studies. The electrotactile stimuli were composed of a continuous sinusoidal carrier with a frequency of 175Hz, which fell within the sensitive range of Pacinian mechanoreceptors (Kandel et al., 2000; Verrillo, 1963) and differed from the auditory tone because we aimed to bias potential audio-tactile interactions toward the auditory noise, not the tone. For the same reason, the carrier of the electrotactile stimulation was amplitude-modulated sinusoidally at the same frequency as the noise (4Hz, depth: 100%, start phase: 0° or 180°; see Figure 1A). The electrotactile stimuli were presented at a tolerable, individually defined current intensity (see Procedure) within the range from 0.8 to 3.0 mA (mean ± SD across participants: 1.9 ± 0.6 mA; peak-to-zero amplitude). This intensity remained fixed throughout the remaining measurements. While the individualization of the stimulation induced inter-individual variability in physical stimulation intensity, it reduced inter-individual variability in perceived intensity, which we deemed crucial for our behavioral and cortical measures. Note that we refer to the electrotactile stimulation as “tactile” in the remainder of this paper for brevity.

#### 2.2.3 Auditory-tactile stimulus presentation

Auditory and tactile stimuli were generated in MATLAB (The MathWorks, Natick, MA) using a sampling rate of 16 kHz and converted simultaneously to analog signals using a multi-channel D/A converter (National Instruments). Auditory stimuli were presented diotically via insert earphones (EARTone 3A) attenuating ambient noise by ≥ 30 dB (SPL). To induce the perception of touch and reduce the impact of movements, tactile stimuli were presented unilaterally above the peripheral median nerve using an isolated bipolar constant-current stimulator (Digitimer DS5, Digitimer, UK) controlled by the D/A converter. Currents were applied to the skin through two square-shaped rubber electrodes (size: 3×3cm^2^) attached to participants’ right forearms using adhesive conductive paste and tape. The electrotactile stimulation did not produce any mechanical vibration or auditory percept. Presentation of the auditory stimulation was delayed by 1ms relative to the tactile stimulation to compensate for potential differences in signal-propagation times and synchronize arrival of the stimuli in the sensory cortices as measured with scalp EEG or magnetoencephalography (Hashimoto et al., 1995; Korvenoja et al., 1999; Picton et al., 1974). Onsets and offsets of all stimulus components were ramped using raised-cosine ramps.

### 2.3 Experimental design

This study used a within-subjects design with two fully crossed factors. The first factor, which we refer to as *Stimulation* (Figure 1B), comprised three levels resembling the different types of stimulation: in-phase audio-tactile stimulation (AT_0_), anti-phase audio-tactile stimulation (AT_180_), and auditory-only stimulation (A, purely auditory stimulation). The second factor, which we refer to as *SNR*, comprised four levels resembling the different auditory SNRs (see Auditory stimuli). The dependent measures were auditory performance (assessed as the percentage of correct responses) and cortical response magnitude or phase (assessed from EEG responses phase-locked to the stimulus-amplitude modulations), which are described further below.

### 2.4 Tasks

Auditory performance was assessed using a two-interval two-alternative forced-choice target detection task. The target was a temporary decrease in the level of the tone, which was implemented by reducing the tone amplitude by 6 dB (SPL) during a 1-s interval (including 10-ms on/off ramps). Each trial began with the presentation of two consecutive, visually-cued 1-s observation intervals (Figure 1C), of which a randomly chosen one contained the target, with equal probability across trials. Participants were instructed to focus their attention on the tone, ignore the noise and the tactile stimulation (if present), and indicate the interval during which they heard the target by pressing one of two buttons with the index finger of their left hand after the second interval. We presumed that the dips in the auditory noise would facilitate listeners in hearing out the tone from the noise and thereby facilitate their ability to detect the target. Presentation of visual stimuli and recording of behavioral responses was implemented in Presentation 20.0 (Neurobehavioral Systems, Inc., Albany CA) and controlled with trigger pulses sent from the D/A converter.

Listeners’ ability to discriminate the two audio-tactile stimulation conditions (AT_0_ and AT_180_) was assessed after the main experiment with a one-interval two-alternative forced-choice audio-tactile synchrony-judgement task. Each trial of this task included the presentation of a single audio-tactile stimulus (either AT_0_ or AT_180_, with equal probability across trials) for 5.5 s (including 1s on/off ramps). Listeners were instructed to judge whether the auditory and tactile stimulation were synchronous or alternating. In both tasks visual feedback on response correctness was given after each response.

### 2.5 Cortical data acquisition

EEG was recorded with 31 Ag/AgCl electrodes mounted in an elastic cap and referenced to the left mastoid (A1), using a BrainAmp amplifier (Brain Products, Munich, Germany). The scalp electrodes were placed according to the International 10-20 system: Fz, FCz, Cz, CPz, Pz, POz, FP1/FP2, F3/F4, F7/F8, FC1/FC2, FC5/FC6, C3/C4, T7/T8, CP1/CP2, CP5/CP6, P3/P4, P7/P8, O1/O2, and A2 (right mastoid). The vertical EOG was recorded with an additional electrode placed below the left eye. The recorded EEG signals were filtered using a lowpass analog filter (cut-off frequency: 250 Hz) and digitized using a sampling rate of 1000 Hz.

### 2.6 Procedure

The experimental procedure included the following steps: (1) After participants gave informed consent, they were seated in a sound-attenuated chamber where they were screened using pure-tone audiometry and a questionnaire assessing their eligibility to undergo transcutaneous median-nerve stimulation. (2) The electrotactile stimulation electrodes were attached to the participants’ right forearms. Participants did an adjustment task, in which they were asked to fine-tune the intensity of the tactile stimulation with a step size of 0.1 mA to the highest level that did not induce discomfort. This was repeated five times and the observed intensities were averaged and fixed for the tactile stimuli used in the remainder of the session. (3) Participants practiced the main task during two short blocks of 20 trials each, which involved both audio-tactile stimulation conditions and various SNRs. They did not receive any instruction or training to integrate the auditory and tactile stimuli. (4) The EEG cap was placed on the participants’ heads, the electrodes were filled with conductive paste, and electrode impedances were reduced to below 10 kΩ. (5) Participants performed the main task in twelve consecutive blocks (corresponding to the 3×4 experimental conditions) during which the EEG was recorded. Each block lasted 4 minutes and contained 60 trials of the given condition. In total 720 trials were presented during the EEG experiment. Before each block, the auditory noise and tactile stimulation were ramped up with some delay relative to the tone (ramp durations: tone: 10ms, noise: 3.5s, tactile stimulation, if presented: 5s) so that participants could immediately hear and track the tone. The stimulation was kept on throughout the entire block (including response and feedback intervals) to avoid neural onset/offset responses and promote a steady state in participants. The order of the twelve blocks was counterbalanced across participants to avoid confounding by potential time-related effects, such as adaptation of tactile-evoked responses. Participants were offered to take short breaks between blocks. (6) After the EEG experiment, participants performed 30 randomly ordered trials (15 trials in condition AT_0_ and 15 trials in condition AT_180_) of the audio-tactile synchrony-judgement task.

### 2.7 Data analysis

#### 2.7.1 EEG data preprocessing

EEG data were processed in MATLAB (2020a, The MathWorks, Inc., Natick, MA) using EEGLAB 14.1.2 (Delorme & Makeig, 2004) and custom scripts. The EEG channel data were re-referenced to the average of all scalp electrodes. To reduce artefacts, independent component analysis was applied using a second-order blind-identification algorithm (Belouchrani et al., 1997). For this analysis the continuous channel data were band-pass filtered using a linear-phase finite-impulse response filter (cutoff frequencies: 1 Hz and 40 Hz, zero phase shift, filter order: 3300). Artifactual components resembling blinks and eye or head movements were identified by visual inspection and rejected (mean ± SD across participants: 6.4 ± 3.1% of components). The weights of the non-artifactual components were reapplied to the original unfiltered channel data, which were subsequently band-pass filtered as above, but using wider cutoff frequencies of 0.5 Hz and 45 Hz (Jaeger et al., 2018). The upper cutoff frequency ensured the removal of line noise and any electrotactile stimulation-induced artefacts, which were limited to a band spanning the carrier frequency of the tactile stimulation ± its amplitude-modulation rate (175 ± 4Hz).

The artefact-reduced continuous channel data were segmented into trials lasting from 50ms before the first observation interval until the end of the second observation interval. A baseline corresponding to the average amplitude between -50 and 0 ms (time-locked to the onset of the first observation interval) was subtracted from each trial. Baseline-corrected trials were further split into 1-s long epochs resembling the two observation intervals, which were labelled as no-target epochs and target epochs according to the presentation of the target. Epochs during which EEG amplitude exceeded ± 75µV at any electrode were deemed to contain residual artefacts and rejected (mean ± SD across participants: 2.5 ± 1.9 % of epochs). On average, 1404 ± 28 epochs per participant were retained for further analysis.

#### 2.7.2 SSR magnitude

The steady-state response (SSR), also called steady-state evoked potential (SSEP), is a rhythmic brain potential evoked by repetitive sensory stimuli, such as the auditory SSR (ASSR), steady-state visually evoked potentials (SSVEP), and SSSEP (steady-state somatosensory evoked potentials) (Norcia et al., 2015; Noss et al., 1996; Picton et al., 2003). SSRs provide a prominent and reliable measure of neural responses phase-locked to the repetition frequency of the sensory input (Breitwieser et al., 2012; Lins & Picton, 1995). Here, we used SSRs to measure the magnitude of phase-locked cortical responses to the different tagging frequencies (noise-modulation rate: 4Hz, tone-modulation rate: 37Hz) in the sensory input.

As the start phase of all amplitude modulations was constant across all blocks, trials, and observation intervals within a given condition, epochs could be averaged across observation intervals in the time domain and were then submitted to a discrete Fourier transform (1000 points, resulting in a spectral resolution of 1Hz). The SSR was calculated by dividing power at the tagging frequency by the noise floor (calculated as the average power of the two frequency bins directly neighbouring the tagging frequency), expressed in units of dB. This was done separately for each EEG channel, condition, and participant.

To focus the neural analyses on cortical responses to the auditory noise and the tone, SSRs were spatially filtered. Given the limited number of EEG channels, we refrained from applying a source-analysis approach (e.g. dipole or distributed source models). Instead, we weighted the SSR at a given EEG channel by the auditory SSR observed at that channel during purely auditory stimulation (condition A); this was done individually for each participant. More specifically, the channel weights (auditory SSRs) were normalized to the range from 0 to 1 at the individual level, multiplied by the participant’s single-channel SSRs observed in another given experimental condition, and the resulting weighted SSRs were averaged across channels by computing their arithmetic mean divided by the sum of the normalized channel weights. For further details on this averaging approach and application of alternative channel-selection approaches to our data, see the Supplementary materials.

For the 4-Hz SSR, the weights were derived from the 4-Hz auditory SSR obtained in condition A at the lowest SNR, thereby focusing the analysis on the cortical representation of the noise-amplitude modulation. Analogously, for the 37-Hz SSR, the weights were derived from the 37-Hz auditory SSR obtained in condition A at the highest SNR, thereby focusing the analysis on the cortical representation of the tone-amplitude modulation. To avoid non-dependency in the data analysis, the aforementioned lowest-SNR condition A and highest-SNR condition A were excluded from subsequent statistical analyses of 4-Hz SSR and 37-Hz SSR, respectively.

#### 2.7.3 SSR Phase

To investigate whether cortical responses to the tactile stimulation coincided with cortical responses to the peaks or dips in the auditory noise, the phase of responses to the 4-Hz modulations was assessed by extracting the phase angle from the corresponding Fourier coefficient. For the multisensory stimulation (condition AT_0_ or AT_180_), SSR phase was extracted from the highest-SNR condition (which was assumed to be dominated by responses to the tactile stimulation), whereas for the unisensory auditory stimulation (condition A), it was extracted from the lowest-SNR condition (which was assumed to be dominated by responses to the auditory noise). According to our hypothesis, stronger phase similarity should be observed between condition A vs. condition AT_0_ than condition A vs. condition AT_180_. To test this, SSR phase was extracted from the channel that responded most strongly to the noise (i.e., strongest channel weight; see above, section SSR magnitude) for each participant. This was done to ensure good signal strength, which can be considered as a prerequisite for obtaining a reliable phase estimate. The extracted phase angles were compared across conditions. All phase calculations were performed in Matlab using the Circular Statistics toolbox (Berens, 2009).

### 2.8 Statistical testing

The single-subject estimates were submitted to second-level (random-effect) statistical analyses. This involved two-way repeated-measures analyses of variance (ANOVAs) including the factors *Stimulation* (two levels resembling conditions: A and AT_0_ or AT_180_) and *SNR* (three levels resembling the different SNRs). Observations more than three SD away from the distribution mean were defined as outliers and discarded. The assumption of normality was verified with Kolmogorov-Smirnov tests, which did not detect any significant deviation from normality (all *P* > 0.05). The assumption of sphericity was assessed with Mauchly’s tests and Greenhouse-Geisser correction was applied to adjust the degrees of freedom when the assumption was violated. Effect sizes were quantified using partial eta-squared (*η*^*2*^_*p*_). Effects of audio-tactile stimulation relative to purely auditory stimulation at individual SNRs were assessed using paired t-tests. Statistical significance of phase difference between conditions (A and AT_0_ or AT_180_) was tested using a Watson-Williams multi-sample test (analogous to a one-way ANOVA). Linear correlation and its significance were assessed using Pearson’s correlation coefficient *r*. A significance criterion *α* = 0.05 was used for all tests. Multiple-comparison correction was done using the false discovery rate (FDR) (Benjamini & Yekutieli, 2001).

## 3. Results

### 3.1 Cortical 4-Hz response magnitude

To investigate the effect of continuous tactile stimulation on cortical responses to auditory noise, we first extracted the cortical representation of the noise. The noise representation was defined as the EEG channels showing the strongest phase-locked responses to the amplitude-modulation of the noise at the highest level in the absence of tactile stimulation (condition A, lowest SNR; see Methods). This resulted in a spatial filter (i.e., topographic map of channel weights) for each participant. The average filter (arithmetic mean across participants) is illustrated in Figure 2A and reported in numerical form in Table S1 in the Supplementary material. The noise representation was on average most prominent over frontocentral cortical regions, especially in the left hemisphere despite the diotic nature of the auditory stimulation. However, statistical comparison of channel weights at corresponding locations in the left vs. right hemisphere provided no evidence for a functional lateralization at any scalp location (|*t* (23)| < 3.26, *p* > .12).

**Figure 2.**
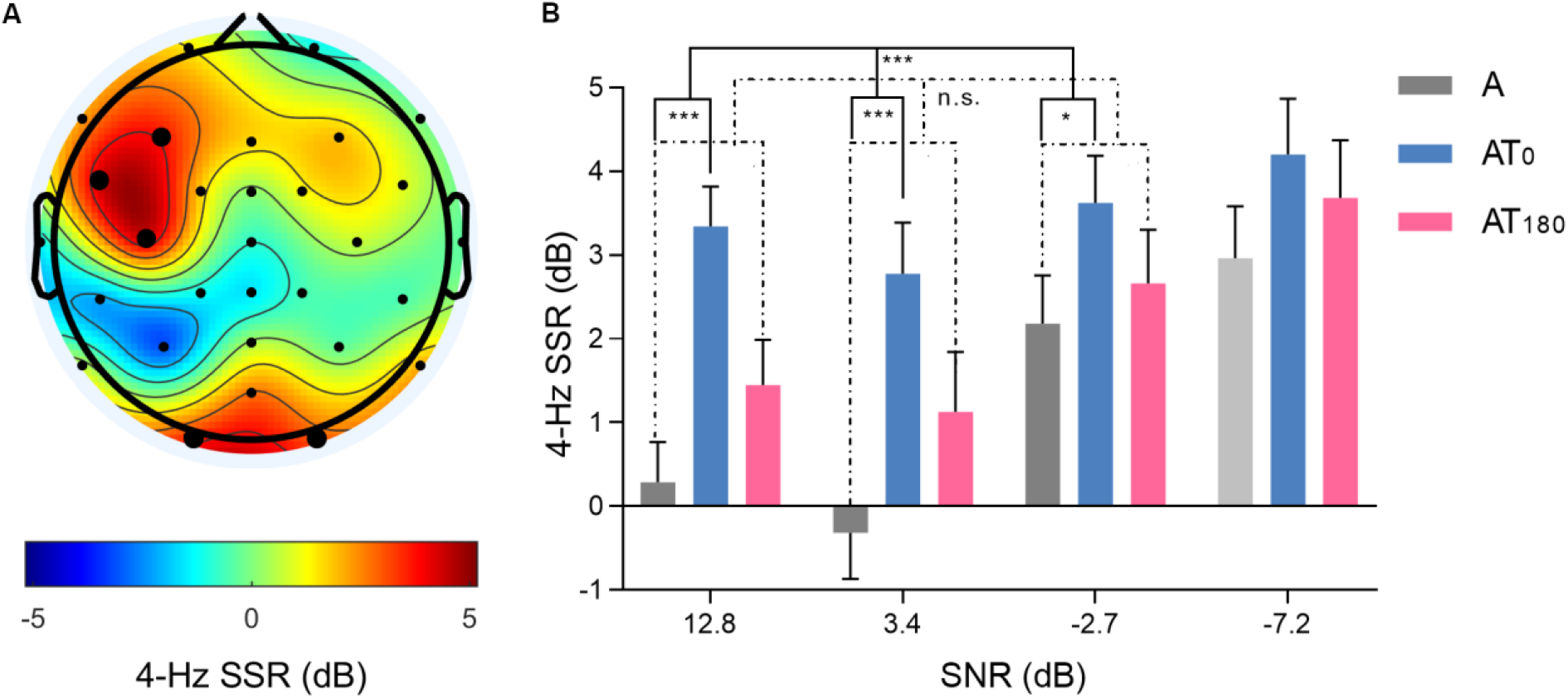
Cortical representation of the amplitude modulation of the auditory noise and average 4-Hz SSR in each condition. **A**. The topographic map shows the average 4-Hz auditory SSR in the purely auditory condition at the lowest SNR. This map was deemed to reflect the cortical representation of the amplitude modulation of the auditory noise and used as a spatial filter to focus the main analysis for tactile (phase) effects on that representation. Data show the mean across participants. Black dots represent scalp locations of the EEG electrodes. Enlarged dots represent locations at which the 4-Hz auditory SSR was significantly above zero (p < 0.05, FDR-corrected). **B**. The bar plot shows the weighted-channel average 4-Hz SSR for each stimulation condition as a function of SNR. The gray bars, blue bars, and magenta bars show respectively data from the purely auditory condition (A), in-phase audio-tactile condition (AT_0_) and anti-phase audio-tactile condition (AT_180_). Condition A at the lowest SNR is shown in light gray as this data point may be biased due to the fact that it defined the spatial filter (panel A). Data show the mean ± SE across participants. Solid lines and dashed lines refer respectively to the main effect of Stimulation for A vs. AT_0_ and A vs. AT_180_. Below the lines, *, *** and n.s. represent p < 0.05, p < 0.001 and no significant difference, respectively.

Figure 2B shows the weighted-channel average 4-Hz SSR for each condition. As expected, responses to the auditory noise were weak or absent when the level of the noise was low, and stronger when the noise level was high. Responses to audio-tactile stimulation were generally stronger than responses to purely auditory stimulation, especially at low noise levels, suggesting that these responses were partially driven by tactile input. Responses to the in-phase tactile stimulation were generally stronger than responses to anti-phase tactile stimulation.

The observation of the strongest responses in the condition comprising in-phase audio-tactile stimulation is conform with our predictions and suggests that in-phase tactile stimulation strengthened cortical responses to the fluctuations of the auditory noise (whether this strengthening reflects multisensory integration is discussed below, see section 4.3). This notion was statistically supported by a two-way repeated-measures ANOVA with factors *Stimulation* (A vs. AT_0_) and *SNR* (12.8 dB, 3.4 dB, and -2.7 dB; the lowest-SNR condition was excluded from this analysis to avoid statistical bias, see Methods) that revealed main effects of *Stimulation* (*F* (1,23) = 22.14, *p* < .001, *η*_*p*_^*2*^ = 0.49) and *SNR* (*F* (2,46) = 7.09, *p* = .002, *η*_*p*_^*2*^ = 0.24), and a significant interaction between the above factors (*F* (2,46) = 3.74, *p* = .031, *η*_*p*_^*2*^ = 0.14). Paired-sample t-tests further showed that the strengthening effect of in-phase tactile stimulation was strongest at the softest noise level (12.8 dB: *t* (23) = 5.48, *p* < .001; 3.4 dB: *t* (23) = 4.21, *p* < .001; -2.7 dB: *t* (23) = 2.05, *p* = .026).

To disambiguate whether this strengthening effect of in-phase tactile stimulation was caused by the tactile stimulation itself or more specifically by its synchrony with the auditory noise, an analogous analysis was applied to the anti-phase condition. A two-way repeated-measures ANOVA with factors *Stimulation* (A vs. AT_180_) and *SNR* (same as above) revealed a main effect of *SNR* as above (*F* (2,46) = 11.09, *p* < .001, *η*_*p*_^*2*^ = 0.33), but no significant main effect of *Stimulation* (*F* (1,23) = 3.22, *p* = .086, *η*_*p*_^*2*^ = 0.12) or interaction (*F* (2,46) = 0.84, *p* = .437, *η*_*p*_^*2*^ = 0.04). This indicates that the strengthening effect of in-phase tactile stimulation as observed above was caused by the audio-tactile phase (i.e., the synchrony of the tactile stimulation with the auditory noise) rather than by the mere presence of the tactile stimulation alone.

### 3.2 Cortical 4-Hz response phase

To verify whether cortical responses to in-phase tactile stimulation were indeed synchronous with cortical responses to the auditory noise, we assessed the phase of 4-Hz EEG responses and tested it for differences between the purely auditory and audio-tactile conditions. To obtain a reliable estimate of cortical phase from each participant, we extracted the 4-Hz phase angle from the individual channel that responded most strongly to the noise, i.e., the channel showing the strongest 4-Hz SSR in the purely auditory condition at the lowest SNR (see Fig. 2A for average channel weights). The average 4-Hz ASSR magnitude of the selected channels was 10.05 ± 3.76 dB (mean ± SD across participants; range: 4.11 dB to 18.61 dB).

The cortical phase-angle distributions extracted from these channels are shown for each relevant condition in Figure 3. The average cortical phase angles were as follows: 282.2 ± 74.5° in condition A at the lowest SNR, 233.5 ± 79.8° in condition AT_0_ at the highest SNR, and 119.7 ± 65.0° in condition AT_180_ at the highest SNR (circular mean ± SD across participants). The average phase difference between condition A vs. condition AT_0_ was 303.1 ± 50.0°, and between condition A vs. condition AT_180_, it was 252.3 ± 72.5° (circular mean ± SD across participants). Statistical comparison of cortical phase in condition A vs. condition AT_0_ revealed no significant difference (*F* (1,46) = 0.67, *p* = .419), suggesting that cortical responses to the auditory noise and cortical responses to in-phase tactile stimulation were not reliably asynchronous. Conversely, the comparison of condition A vs. condition AT_180_ revealed a main effect of *Stimulation* (*F* (1,46) = 15.76, *p* < .001), indicating that cortical responses to the auditory noise and cortical responses to anti-phase tactile stimulation were asynchronous as expected.

**Figure 3.**
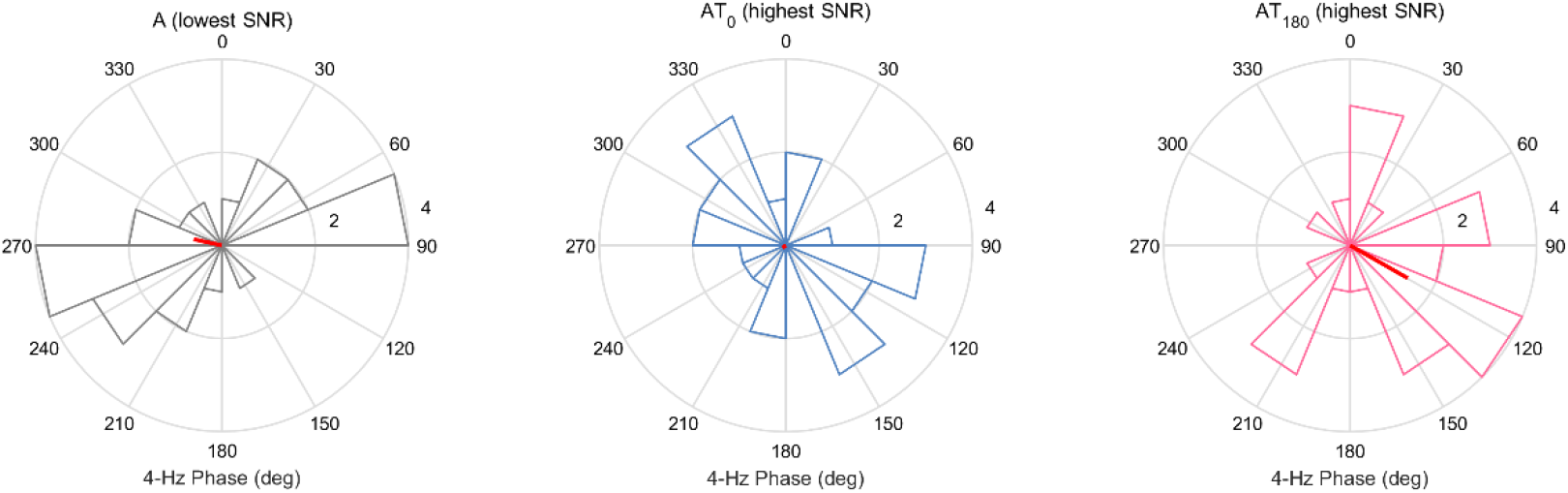
Phase of 4-Hz EEG responses. The polar histograms show 4-Hz phase for the EEG channel that responded most strongly to the noise (i.e., the individual channel showing the maximum 4-Hz SSR in condition A at the lowest SNR). From left to right are shown the purely auditory condition (condition A), in-phase audio-tactile condition (condition AT_0_), and anti-phase audio-tactile condition (condition AT_180_). Red lines represent the mean angle (direction of the red line) and consistency (length of the red line) of 4-Hz phase across all participants. The circle radius represents the count of participants.

### 3.3 Cortical 37-Hz response magnitude

To investigate the effect of continuous tactile stimulation on cortical responses to the tone, we first extracted the cortical representation of the tone, analogous to the 4-Hz analyses above. The cortical representation of the tone was defined as the EEG channels showing the strongest phase-locked responses to the 37-Hz amplitude-modulation of the tone, given the softest noise level in the absence of tactile stimulation. The resulting map of channel weights (37-Hz auditory SSRs) is shown in Figure 4A (for a numerical representation, see Supplementary Table 1). The tone representation was most prominent over fronto-central cortical regions, consistent with previous auditory SSR results (Herdman et al., 2002; Schwarz & Taylor, 2005).

**Figure 4.**
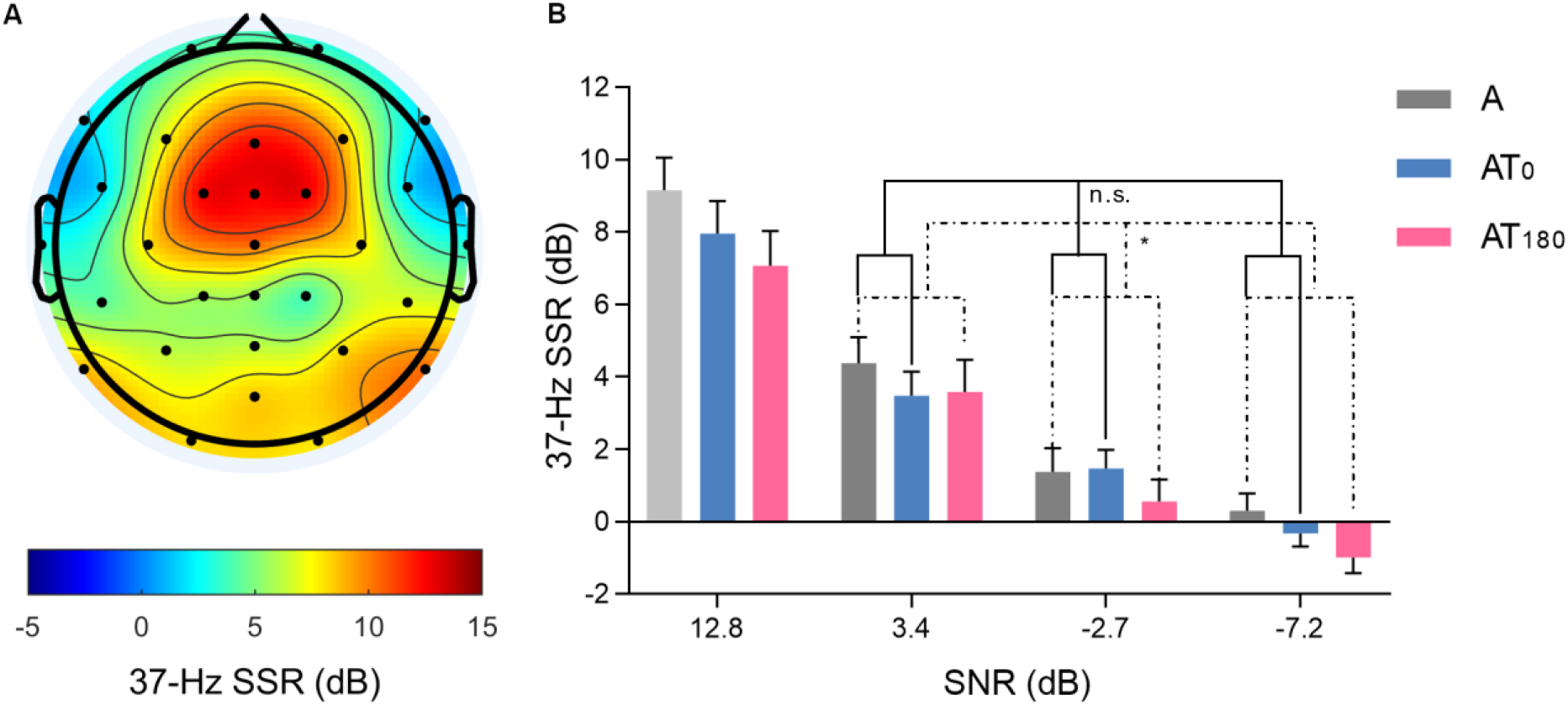
Cortical representation of the amplitude modulation of the tone and average 37-Hz auditory SSR. **A**. The topographic map shows the average 37-Hz auditory SSR in the purely auditory condition at the highest SNR. This map was deemed to reflect the cortical representation of the amplitude modulation of the tone and used as a spatial filter to focus the main analysis for tactile (phase) effects on that representation. Data show the mean across participants. **B**. The bar plot shows the weighted-channel average 37-Hz auditory SSR for each stimulation condition as a function of SNR. Color coding is as in Figure 2B. Condition A at the highest SNR is shown in light gray as this data point may be biased by the fact that it defined the spatial filter (panel A). Data show the mean ± SE across participants. Below the lines, *, ** and n.s. represent p < 0.05, p < 0.01 and no significant difference, respectively.

Figure 4B shows the weighted-channel average 37-Hz auditory SSR for each condition. As expected, responses to the tone were strong when the level of the noise was low, and weaker or absent when the noise level was high. Responses to audio-tactile stimulation were on average slightly weaker than responses to purely auditory stimulation. Responses to the anti-phase tactile stimulation were on average weaker than responses to in-phase tactile stimulation.

The observation of the weakest responses in the condition comprising anti-phase audio-tactile stimulation is conform with our predictions and suggests that anti-phase tactile stimulation masked cortical responses to the tone during the dips in the auditory noise. This notion was statistically supported by a two-way repeated-measures ANOVA with factors *Stimulation* (A vs. AT_180_) and *SNR* (3.4 dB, -2.7 dB, and -7.2 dB; the highest-SNR condition was excluded from this analysis to avoid statistical bias, see Methods), which revealed main effects of *Stimulation* (*F* (1,23) = 6.58, *p* = .017, *η*_*p2*_ = 0.22) and *SNR* (*F* (1.4,32.1) = 38.28, *p* < .001, *η*_*p*_^*2*^ = 0.63), and no significant interaction (*F* (2,46) = 0.30, *p* = .744, *η*_*p*_^*2*^ = 0.03).

To disambiguate whether the neural masking effect of anti-phase tactile stimulation on the tone representation was caused by the tactile stimulation itself or more specifically by its synchrony with the dips in the auditory noise, an analogous analysis was applied to the in-phase audio-tactile condition. A two-way repeated-measures ANOVA with factors *Stimulation* (A vs. AT_0_) and *SNR* (same as above) revealed a main effect of *SNR* as above (*F* (2,46) = 26.82, *p* < .001, *η*_*p*_^*2*^ = 0.71), but no significant main effect of *Stimulation* (*F* (1,23) = 1.92, *p* = .180, *η*_*p*_^*2*^ = 0.08) or interaction (*F* (2,46) = 1.68, *p* = .209, *η*_*p*_^*2*^ = 0.13). Thus, in-phase tactile stimulation did not significantly enhance tone-evoked responses, contradicting our original hypothesis that in-phase tactile stimulation would enhance dip listening to the tone. The effect of anti-phase tactile stimulation on tone-evoked responses and the null effect of in-phase tactile stimulation together indicate that the former effect was caused by the audio-tactile phase (i.e., the synchrony of the anti-phase tactile stimulation and the dips in the auditory noise) rather than by the mere presence of the tactile stimulation alone.

### 3.4 Auditory performance

To further investigate the effect of tactile stimulation on dip listening at the behavioral level, we assessed listeners’ perception of the tone based on their auditory target-detection performance. The results are shown for each condition in Figure 5. Overall, listeners’ performance ranged on average from 90.8 ± 1.5 % at the highest SNR to 55.3 ± 1.3 % at the lowest SNR (chance level: 50%). Contrary to our predictions, we observed no systematic difference in auditory performance across the purely auditory vs. audio-tactile conditions: two-way repeated-measures ANOVAs with factors *Stimulation* (A and AT_0_, or A and AT_180_) and *SNR* (12.8 dB, 3.4 dB, -2.7 dB, and -7.2 dB) revealed a main effect of *SNR* as expected, but no main effect of *Stimulation* and no significant interaction, regardless of the phase of the tactile stimulation (AT_0_: *SNR*: *F* (3,69) = 178.10, *p* < .001, *η*_*p2*_ = 0.89, *Stimulation*: *F* (1,23) = 0.22, *p* = .645, *η*_*p2*_ = 0.01, *interaction*: *F* (3,69) = 0.40, *p* = .754, *η*_*p2*_ = 0.02; AT_180_: *SNR*: *F* (3,69) = 146.67, *p* < .001, *η*_*p2*_ = 0.86, *Stimulation*: *F* (1,23) = 0.12, *p* = .736, *η*_*p2*_ = 0.005, *interaction*: *F* (3,69) = 1.32, *p* = .275, *η*_*p2*_ = 0.05). Focusing the behavioural analysis on the SNR conditions revealing the strongest neural effects (in-phase stimulation: second-highest SNR; anti-phase stimulation: lowest SNR) did not qualitatively change this outcome (AT_0_ > A at 3.4 dB: *t* (23) = 0.34, *p* = .370; AT_180_ < A at -7.2 dB: *t* (23) = 1.26, *p* = .110). Thus, in contrast to listeners’ cortical responses, their auditory performance was not significantly affected by the tactile stimulation or its relative phase.

**Figure 5.**
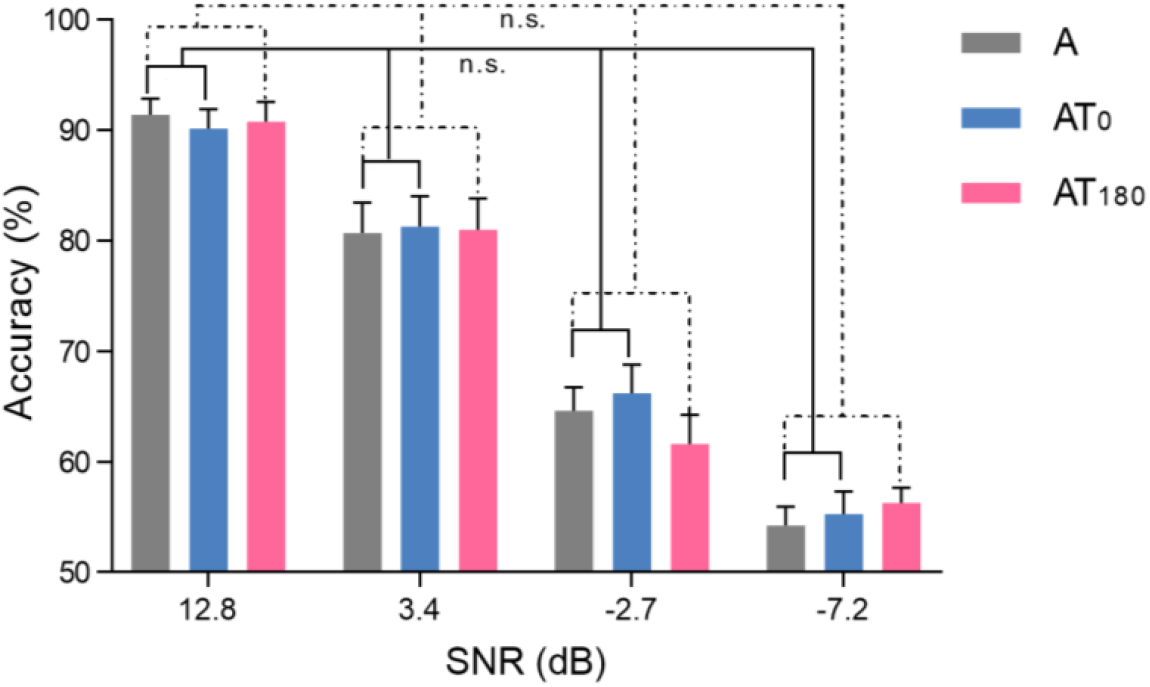
Behavioral performance on the auditory target-detection task. The bar plot shows listeners’ average response accuracy for each stimulation condition as a function of SNR. The chance level was 50%. Color coding is as in Figure 2B. Data show the mean ± SE across participants. Below the lines, n.s. represents no significant difference.

### 3.5 Audio-tactile synchrony judgments

In the main experiment, listeners were instructed to ignore the tactile stimulation (see Methods), leaving unclear whether paying attention to the tactile stimulation would have enabled them to identify and possibly exploit the audio-tactile phase. We explored this in a subsequent behavioral test using an audio-tactile synchrony-judgment task, which assessed participants’ ability to directly discriminate between the in-phase and anti-phase audio-tactile stimulation (AT_0_ vs. AT_180_). We found that participants performed this task on average at a level of 49.9 ± 11.5% (mean ± SD; range: 33.3% to 76.7%), which did not differ significantly from the chance level of 50% (one-tailed paired t-test: *t* (23) = 0.06, *p* = .953). This indicates that participants were unable to reliably identify the relative phase of the auditory and tactile stimuli even when they intentionally paid attention to the tactile stimulation.

## 4. Discussion

In the present study, we tested whether long-lasting tactile input that fluctuates in phase or anti-phase with an ongoing auditory noise can respectively amplify the fluctuations of the noise or mask the dips in the noise. We found that in-phase tactile stimulation (compared with purely auditory stimulation) increases cortical responses to the fluctuations of the auditory noise, as measured with EEG on the scalp. We further found that anti-phase tactile stimulation (compared with purely auditory stimulation) decreases cortical responses to a tone presented during the noise dips. In contrast to these cortical effects, we observed no behavioral effect of tactile stimulation or its relative phase on auditory target-detection performance, which served here as a measure of dip listening. In sum, these results show that the relative phase of ongoing rhythmic audio-tactile stimulation can influence cortical responses to fluctuating auditory noise and a tone embedded in that noise, without a corresponding impact on dip listening to the tone.

### 4.1 In-phase tactile stimulation enhances cortical responses to fluctuating auditory noise

Our first observation shows that in-phase tactile stimulation (compared with purely auditory stimulation) can increase the 4-Hz SSR, a measure quantifying here the magnitude of phase-locked cortical responses to the fluctuations of the auditory and tactile stimulation. The strengthening effect was not observed when comparing anti-phase tactile stimulation with purely auditory stimulation, indicating that the effect was caused by the audio-tactile synchrony, not the tactile stimulation itself. The enhancing effect of synchronous tactile stimulation is consistent with previous neuroimaging results (Foxe et al., 2002; Hoefer et al., 2013; Murray et al., 2005) and parallels behavioral results showing auditory enhancement by brief synchronous tactile inputs (Gillmeister & Eimer, 2007; Schürmann et al., 2004; Wilson et al., 2009, 2010). The pattern of results suggests that cortical responses to continuous fluctuations of auditory noise and tactile stimulation as measured on the scalp add up: in-phase stimulation results in an overall increased response, whereas anti-phase stimulation produces no significant change. Whether the auditory and tactile responses add linearly or integrate in a non-linear manner is discussed below (see section 4.3).

Our interpretation is partially supported by the 4-Hz phase results: we observed that anti-phase tactile stimulation, but not in-phase tactile stimulation, induced a significant phase shift in cortical responses to the auditory noise. This suggests that the audio-tactile phase differences in the physical stimuli were partially preserved in the cortex, i.e., cortical responses to the peaks in the in-phase tactile input coincided approximately with responses to the peaks in the auditory noise, whereas responses to the peaks in the anti-phase tactile input coincided approximately with responses to the auditory noise dips. This interpretation is in line with previous audio-tactile EEG results based on faster modulation rates of up to 40-Hz (Timora & Budd, 2013, 2018) and the more general notion that tactile inputs can enhance cortical responses to auditory stimuli (see Introduction). It mismatches with the observation of no auditory effect of audio-tactile phase in a previous study that used unmodulated 250-Hz sinusoidal tactile stimulation (Wilson et al., 2009), indicating that cortical audio-tactile phase effects may emerge primarily at low (≤ 40Hz) modulation rates. Overall this suggests that ongoing tactile input that slowly fluctuates in phase with an auditory noise can enhance the fluctuations of that noise in the cortex in a sustained manner. Put differently, the previously observed enhancing effect of brief tactile inputs on auditory-evoked responses can operate in a slow continuous manner, and the relative phase of the ongoing tactile input appears to play an important role in this.

Despite our observation of an effect of audio-tactile phase on auditory noise-evoked responses, our participants were essentially unable to correctly identify that phase, even when they intentionally paid attention to it, as suggested by our audio-tactile synchrony-judgment results. This discrepancy between neural and behavioral results indicates that audio-tactile phase exhibited salient bottom-up effects on cortical responses without the participants’ awareness of this phase.

Our interpretation of somatosensorily-induced enhancement of auditory-evoked cortical responses needs to be treated with some caution for two reasons. Firstly, by “cortical representation” we refer to neuroelectric signals measured on the scalp, which cannot be attributed specifically to auditory vs somatosensory cortical sources due to volume conduction (Ozcan et al., 2005; Rodgers et al., 2008). Secondly, the phase of these scalp signals depends on a variety of factors, including the number and relative orientation of the underlying neural sources and potential non-linear mixing of their signals (Nunez et al., 1997), which cannot be easily disentangled here given the limited number of EEG channels. Although we designed our methods to emphasize auditory cortical processes, we cannot exclude partial contributions from tactile processes, as suggested by the observation of strong cortical responses to audio-tactile stimulation at low noise levels. However, we observed comparatively strong cortical effects of noise level, suggesting that our results reflect predominantly auditory processes. Future studies of audio-tactile neural integration may address these limitations and separate auditory and tactile processes more effectively with methods offering superior spatial specificity, such as invasive intracranial electrophysiological recordings.

### 4.2 Anti-phase tactile stimulation masks cortical responses to a tone presented during dips in fluctuating noise without a concomitant auditory effect

We observed that the 37-Hz SSR, a measure quantifying here the magnitude of phase-locked cortical responses to the fluctuations of a continuous tone in noise, increases as a function of SNR. Given that the tone level and its modulation remained unchanged in our study, this observation suggests that the 37-Hz SSR reflected the audibility of the tone in the noise rather than the tone level itself, a notion that is in line with previous findings (Billings et al., 2009; Christensen et al., 2019; Phillips, 1990; Teschner et al., 2016; Zhou & Wang, 2010). Importantly, we found that anti-phase tactile stimulation (compared with purely auditory stimulation) induced a decrease in this measure. This suppressive effect was not observed when comparing in-phase tactile stimulation with purely auditory stimulation, indicating that the effect was caused by the anti-phasic relation of the audio-tactile stimulation, not the tactile stimulation itself. Together with our phase results above, which show that cortical responses to anti-phase tactile peaks approximately coincided with responses to the noise dips, this pattern of results suggests that cortical responses to long-lasting tactile input that alternates with fluctuating noise can mask cortical responses to a signal during the noise dips. A possible interpretation of the suppressive effect is that the anti-phase tactile stimulation induced random variation in the cortical processing of the tone, which resulted in reduced phase-locked responses to the tone. Alternatively, the effect may have originated from an attentional modulation of the cortical processing of the tone, induced by potential attentional distraction caused by an increased saliency of the tactile stimulation during the noise dips.

Following our neural observations, we tested for an auditory masking effect, i.e, putative consequences of the anti-phase tactile stimulation on the audibility of the target in the tone. Contrary to our hypothesis, our neural results, and previous findings from brief tactile (Gick, 2008; Gick & Derrick, 2009; Gillmeister & Eimer, 2007; Reed et al., 2021; Reed et al., 2019; Schürmann et al., 2004; Wilson et al., 2009, 2010) or transcranial electric stimulation (Riecke et al., 2015), we found no such auditory effect: neither tactile stimulation nor its relative phase had a significant impact on listeners’ target-detection performance.

One possible explanation for this null result may be that the originally expected auditory enhancement from audio-tactile phase requires listeners to be aware of this phase or pay attention to it. It has been argued that salient tactile onsets/offsets may modulate auditory perception by boosting the listeners’ vigilance (Al-Shargie et al., 2019; Arrabito et al., 2015; Zhang et al., 2016). The 4-Hz periodicity and ongoing nature of our audio-tactile stimulation probably resulted in perceptual integration and/or adaptation to the constituent auditory and tactile pulses, which may have reduced the saliency of individual pulses, resulting in a lack of phase-related variations in vigilance. In fact, our participants were unable to reliably identify the audio-tactile phase, suggesting that they were unaware of it and could not exploit it as a cue. The same may hold for even more rapidly fluctuating audio-tactile stimuli (40Hz) as used by Timora and Budd (2013, 2018), who used longer-lasting stimuli (1.5 s) and similarly found no effect of audio-tactile modulation rates on auditory perception. In their study, the relative amplitude-modulation rate of auditory and tactile stimulation was varied. The results showed no effect of tactile stimulation or its relative modulation rate on listeners’ ability to detect auditory amplitude modulation. Similarly, Wilson et al. (2009) varied the relative phase of unmodulated audio-tactile stimuli with carrier frequencies up to 250Hz and found no effect of tactile stimulation on auditory perception. Tactile onsets/offsets may be more salient when they occur more slowly or unexpectedly, as in the more irregular and less predictable speech-shaped tactile stimuli used in some audio-tactile speech studies (see Introduction). The increased vigilance resulting from such irregular onsets/offsets may further bias decisional processes (Rizza et al., 2018), which may thus explain some of the previously observed positive effects on speech intelligibility.

Another possible explanation for our behavioral null result is that our measure of dip listening may not have been sensitive enough. The auditory task required listeners to discriminate the intensity of the tone across the two intervals (the tone intensity was reduced during the target interval by 6dB; see Methods). We observed a strong effect of noise level on listeners’ performance on this task, indicating that task performance depended on the audibility of the tone in the noise (Hawkins et al., 1950; Miller, 1974). The anti-phase tactile stimulation likely also affected the audibility of the tone, as suggested by our 37-Hz SSR results, and therefore, it should have affected task performance as well. However, this presumed cross-modal effect on tone audibility was probably much weaker than the observed intramodal effect of noise level and therewith insufficient to affect the discriminability of the intensity of the tone.

To further investigate this idea, we conducted additional analyses exploring the effect of audio-tactile phase on the discriminability of the tone’s intensity in the cortex (see Supplementary Materials). We extracted a cortical measure of tone-intensity discriminability (by subtracting the 37-Hz SSR in the target interval from the 37-Hz SSR observed in the no-target interval). We observed that this measure took positive values in all conditions, in line with the observed SNR effect on 37-Hz SSR. However, neither in-phase or anti-phase tactile stimulation (compared with purely auditory stimulation) had a significant effect on this cortical discriminability measure. This observation is in line with the notion that audio-tactile stimulation and its phase influenced the audibility of the tone without altering the discriminability of its intensity.

Future research may investigate auditory masking effects of continuous tactile stimulation with directly task-relevant tactile stimulation that requires participants to identify and exploit the relative phase. In addition, an improved paradigm that provides a more direct measure of signal audibility during the noise dips (e.g., a two-interval forced-choice task, in which one interval contains anti-phase audio-tactile stimulation, the other contains in-phase audio-tactile stimulation, and listeners report the interval during which an auditory signal of fixed intensity appeared louder to them) may prove useful.

### 4.3 A phase effect on multisensory integration

As described above, our results show that scalp-level measures of auditory- and tactile-evoked cortical responses added up in a phase-dependent manner. An important question is whether they added up in a linear or non-linear manner—the latter would hint at integration of the multisensory input (for an overview of multisensory integration criteria, see Stevenson (2014)). We observed that the strengthening effect of in-phase tactile stimulation on cortical responses to the auditory noise (see section 4.1) decreased significantly at louder noise levels. This modulating effect of noise level indicates that responses to auditory and tactile input added up in dependence of the auditory input, rather than in a simple linear manner. The modulating effect of noise level cannot be explained by a potential ceiling effect (where intense auditory input would saturate auditory cortical neurons and thereby leave little headroom for tactile input to further enhance the neurons’ responses): we observed that responses to the in-phase audio-tactile stimulation still increased beyond the loudest noise level that was included in the aforementioned interaction analysis (see Figure 2B, blue rightmost bars).

A more likely explanation for our observation of non-linear summation is an integration of the audio-tactile stimuli. It has been proposed that multisensory integration follows several principles (Holmes & Spence, 2005) and our results seem to conform to them: according to the inverse effectiveness principle, integration of multisensory inputs is more probable and/or effective when responses to the unisensory inputs are relatively weak (Holmes, 2007; Meredith & Stein, 1986; Senkowski et al., 2011). Thus, the in-phase tactile stimulation in our study should have integrated more effectively with the auditory input when the latter was relatively weak (i.e., in the highest-SNR condition) compared to when it was rather intense (i.e., in the lower-SNR conditions). Our results indeed match this expectation by showing that the strongest tactile stimulation effect occurred when the auditory input was weakest (see section 3.1). Moreover, according to the temporal principle, integration of multisensory inputs is more probable and/or effective when responses to the unisensory inputs are more synchronous (Holmes & Spence, 2005). Support for this principle comes from previous studies using audio-tactile stimuli with clear onsets/offset and relative high frequencies (Timora & Budd, 2013, 2018; Wilson et al., 2010). Presuming that the temporal principle can be extended from onsets/offsets to ongoing periodic signals, the tactile stimulation and the auditory input in our study should have integrated more effectively when their evoked responses were more synchronous (i.e., in the in-phase condition) compared to when they were asynchronous (i.e., in the anti-phase condition). Our results match this expectation by showing both an effect of in-phase (but not anti-phase) tactile stimulation on responses evoked by the auditory noise and an effect of anti-phase (but not in-phase) tactile stimulation on responses evoked by the auditory tone. Regarding the latter observation, it should be noted that the tone was most salient during the noise dips and these dips were *synchronous* with the *anti*-phase tactile stimulation. In contrast to the facilitating effect on responses to the noise, the effect on tone-evoked responses was of suppressive nature. This likely reflects the fact that tactile stimulation and tone physically fluctuated at unrelated rates (4Hz vs 37Hz), which likely reduced the synchrony of their amplitude envelope-evoked responses and thereby hampered their neural integration (Budd & Timora, 2013).

In sum, the observed pattern of neural results conforms to principles of multisensory integration of brief audio-tactile events. This suggests that the long-lasting tactile stimulation and synchronous auditory input were integrated in the cortex, despite their lack of perceivable onset cues, which is in line with previous EEG results on continuous audio-tactile speech integration (Riecke et al., 2019). Future studies may further verify this notion by comparing electrophysiologic responses to continuous individual unisensory tactile and auditory stimuli with responses to combined audio-tactile stimuli and applying a super-additivity criterion (Kayser et al., 2005; Sperdin et al., 2009).

A potential mechanism underlying the observed neural integration involves somatosensorily-induced phase shifts in endogenous oscillations in the primary auditory cortex (PAC). Electrophysiological recordings have shown that tactile pulses can adjust the phase of slow endogenous oscillations in monkey PAC to cortical arrival times of auditory inputs and thereby modulate the processing of these inputs (Lakatos et al., 2007). While stimulus-evoked and endogenous oscillations cannot be disentangled in our study, it is likely that the in-phase tactile stimulation supported the alignment of auditory cortical oscillations to the fluctuations in the auditory input and thereby enhanced the processing of this input, whereas the anti-phase tactile stimulation had no such effect.

## 5. Conclusion

Our study shows that ongoing periodic tactile input can influence scalp-level measures of auditory-evoked cortical processing in a sustained manner: tactile input that fluctuates in-phase with auditory noise can enhance the fluctuations of the noise, whereas tactile input that fluctuates in anti-phase with the noise can reduce an auditory signal during the noise dips. Although these effects appear to follow known principles of multisensory integration of brief audio-tactile events in the cortex, they may not necessarily lead to corresponding changes in auditory signal perception, which might require more perceptually salient tactile onsets/offsets and less predictable auditory inputs.

## Acknowledgement

We thank Karlotta Staiger for assistance with data acquisition and Min Wu, Minne Pijfers, Lidongsheng Xing, and three anonymous reviewers for constructive comments on earlier versions of the manuscript. This work was supported by Maastricht University and China Scholarship Council (CSC202008210294 to XF).

## Data availability

The behavioral and EEG datasets for this study are available upon request from the corresponding author.

